# Attenuation of value-to-evidence translation drives biased decision making in anxiety and depression

**DOI:** 10.64898/2026.06.09.731091

**Authors:** Mrugsen Nagsen Gopnarayan, Feng Sheng, Michael Louis Platt, Arjun Ramakrishnan

**Affiliations:** University of Hamburg, Hamburg, Germany; Biological Sciences and Bioengineering Department, Mehta Family Centre for Engineering in Medicine, Indian Institute of Technology Kanpur, Kanpur, India; Neuromanagement Laboratory, School of Management, Zhejiang University, Hangzhou, China; State Key Laboratory of Brain-Machine Intelligence, Zhejiang University, Hangzhou, China; Department of Neuroscience, Perelman School of Medicine, University of Pennsylvania, Philadelphia, PA, USA; Department of Marketing, The Wharton School, University of Pennsylvania, Philadelphia, PA, USA; Department of Psychology, School of Arts and Sciences, University of Pennsylvania, Philadelphia, PA, USA

**Keywords:** anxiety, depression, decision making, drift diffusion model, EEG, centroparietal positivity

## Abstract

Anxiety and depression are globally prevalent conditions associated with maladaptive decision making. However, whether affective symptoms primarily amplify threat avoidance or dampen motivational drive remains debated, and behavioural studies yield inconsistent findings. Here we show that both anxiety and depression impair the fundamental cognitive process of translating objective value into decision evidence. Across independent cohorts from the US and India, participants evaluated risky gambles while we assessed choice behaviour and the centroparietal positivity, an EEG marker of accumulating decision evidence. Prospect theory parameters, like risk and loss aversion, showed little association with symptom severity. Conversely, hierarchical drift–diffusion modelling revealed that higher symptom scores predicted attenuated value sensitivity during evidence accumulation, whereas decision caution remained intact. This reduction in value sensitivity suggests internalizing symptoms disrupt choice at the value-to-evidence interface, offering a unified mechanism underlying biased decision making in affective disorders.

## 1 Introduction

Anxiety and depression are among the most common mental health conditions worldwide, affecting hundreds of millions of people [1]. These disorders exert broad impact on cognition and emotion, particularly decision-making, the process by which individuals assess risks and rewards in uncertain environments [2, 3]. Distortions in decision-making are thought to contribute directly to the persistence of maladaptive behaviours in affective disorders, influencing how individuals respond to potential threats, opportunities, and uncertainty in everyday life [2]. Yet despite the central role of decision processes in anxiety and depression, the mechanisms through which affective symptoms alter the evaluation and use of value information during choice remain poorly understood [4]. Identifying these mechanisms is therefore critical for linking affective symptoms to the cognitive computations that guide behaviour in order to develop better, more targeted interventions.

Risky decision-making is often examined using gambling tasks in which participants choose between options that differ in potential gains and losses. Findings from such paradigms have been frequently interpreted through the lens of *Prospect Theory* [5], which emphasizes two common behavioural biases: loss aversion, in which losses are weighted more strongly than gains of equivalent magnitudes, and risk aversion, characterized by reduced sensitivity to potential gains and losses or increased sensitivity to uncertainty. However, studies linking these constructs to anxiety and depression have yielded inconsistent results [6]. With respect to risk aversion, several studies report increases associated with anxiety [7–9] and depression [10, 11], whereas others find no change in anxiety [12] or even reductions in depression [13]. Similarly, evidence regarding loss aversion is mixed: some work reports increases in anxiety ([12]) and depression ([14]; [15]; [11]), whereas other studies report null effects in anxiety [16] or decreases in depression [17]. These inconsistencies suggest that descriptive models of decision outcomes may be insufficient to capture the mechanisms through which affective symptoms influence decision making.

One reason for this insufficiency is that different theoretical accounts make similar predictions at the level of observable choices while arising from distinct underlying mechanisms. Two broad frameworks have been proposed to explain how anxiety and depression alter decision-making. Threat-avoidance accounts propose that anxiety increases sensitivity to potential losses and promotes cautious responding in the face of uncertainty [2, 18]. In contrast, reduced-motivation accounts emphasize diminished reward responsiveness and reduced approach drive, particularly in depression [19, 20]. Both frameworks can produce superficially similar behavioural patterns in gambling tasks. Although gambling tasks are not clinical assessments, they formalize core features of affective decision making, including uncertainty, risk, reward, and potential loss. Thus, similar behavioural patterns—such as reduced willingness to accept risky options—making them difficult to disentangle using descriptive models alone, motivating the need for neurobiologically-grounded computational frameworks.

Sequential sampling models, such as the Drift Diffusion Model (DDM; Fig. 2a), provide a complementary, chronometric approach by decomposing the decision process [21]. Within this framework, decisions arise through the gradual accumulation of evidence to one of two decision boundaries. The drift rate (v) reflects the strength with which value information drives the decision variable, the decision threshold (a) reflects the level of evidence required before committing to a choice, and the starting point (z) captures any pre-existing bias toward one option. In gambling tasks, drift rate can be decomposed into contributions from potential gains and losses, enabling the estimation of gain drift and loss drift parameters that reflect the influence of reward and punishment on the decision process [22, 23]. In this framework, value signals influence behaviour by shaping the evidence that drives accumulation—a stage we refer to as the value-to-evidence interface. This decomposition allows value sensitivity to be separated from decision caution and bias, offering a compact mathematical framework for distinguishing motivational and control processes that may be differentially altered in anxiety and depression.

**Fig. 1.**
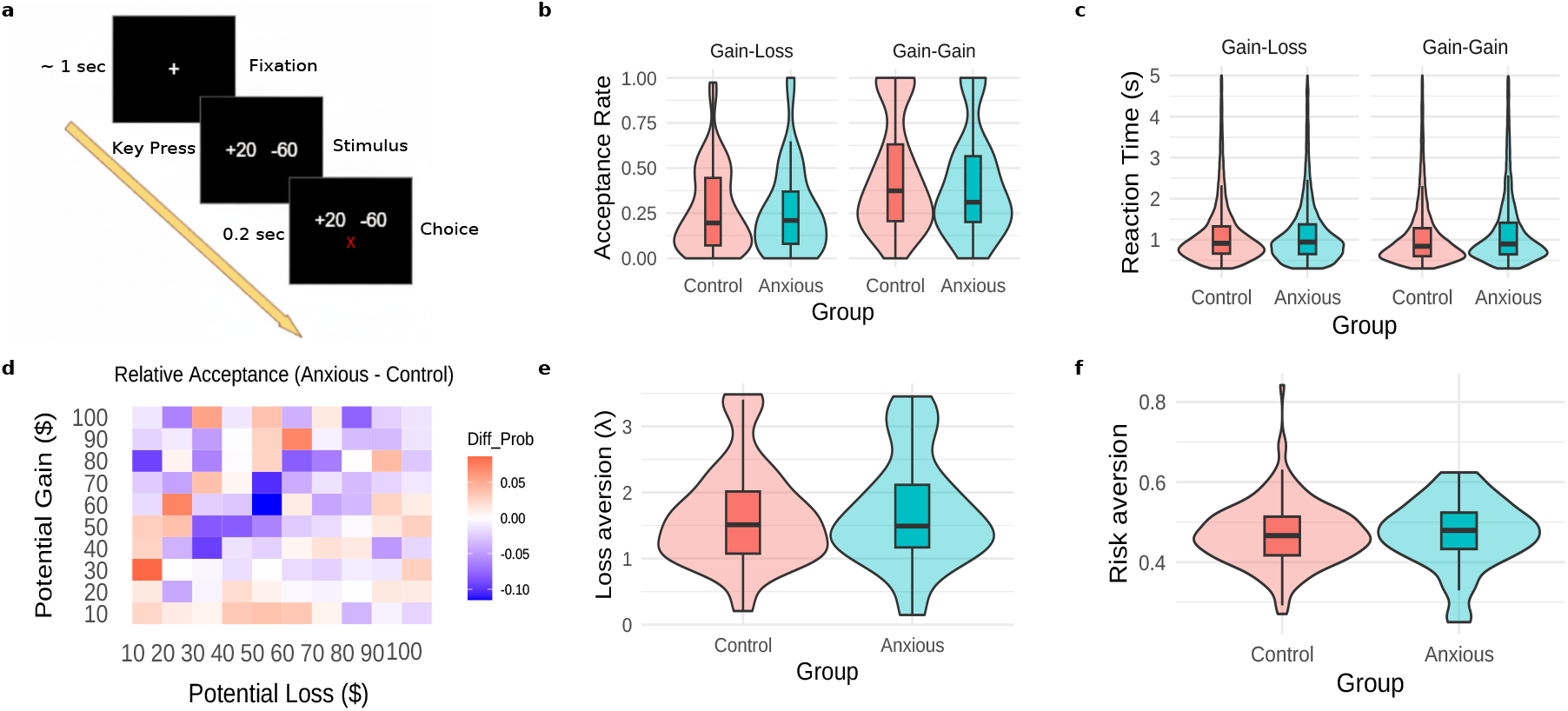
Gambling task and behavioural profiles for anxiety and control groups. **a**, Schematic illustration of the gambling task in which participants repeatedly decide whether to accept or reject mixed gambles offering potential monetary gains and losses in the Gain-Loss condition or potential high gain and low gain against a fixed reward of 100 in Gain-Gain condition. **b-c**, Behavioural outcomes in control versus anxious participants. Panel **b**, shows acceptance rates as a function of gamble characteristics, and **c**, corresponding response times. Both acceptance rate and response times show no significant difference between the two groups. **d**, Heat map of difference in acceptance rate for the two groups for all possible gambles. **e-f**, Prospect theory-based estimates of loss and risk aversion parameters derived from individual choice patterns. Both anxious and control groups show similar parameter distributions.

**Fig. 2.**
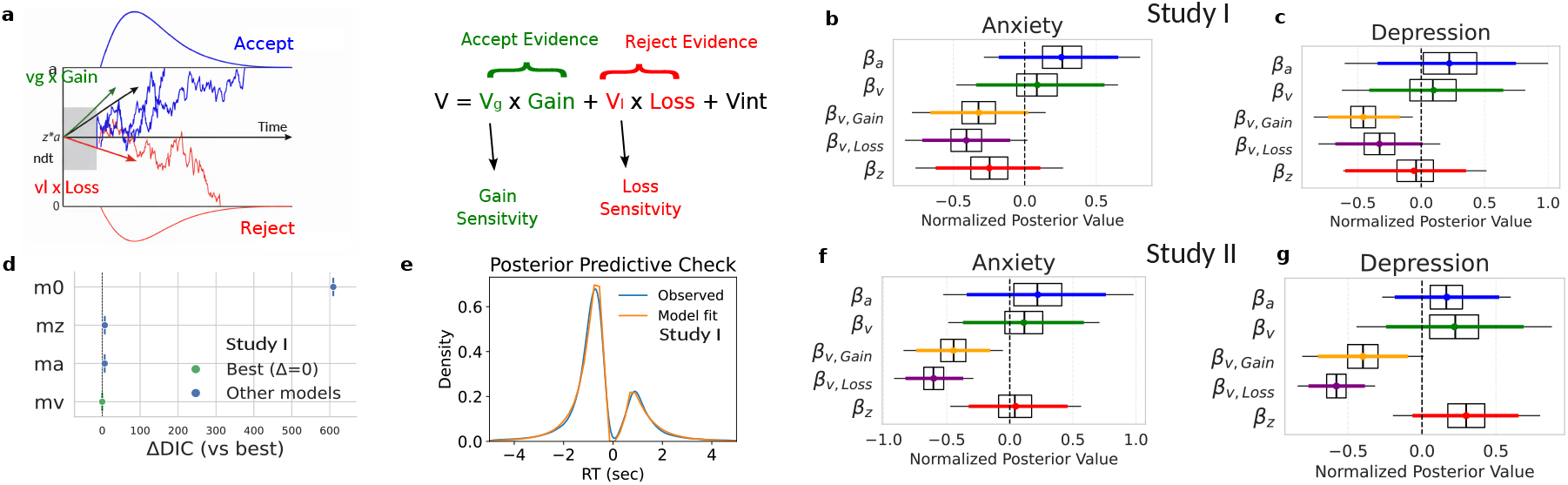
Drift diffusion modelling of Anxiety and Depression in the Gain-Loss condition. **a**, Schematic of the drift diffusion model (DDM). Decisions are modelled as noisy evidence accumulation toward an accept (upper) or reject (lower) boundary. Drift rate is defined as *v* = *v*_*G*_ ×*Gain* −*v*_*L*_ ×*Loss* + *v*_*int*_, where *v*_*G*_ and *v*_*L*_ index gain and loss sensitivity. **b-c**, Posterior distributions for regression coefficients on DDM parameters in Study I (Gain-Loss condition) as a function of continuous, z-score-normalized anxiety and depression scores. Each plot shows standardized effects on boundary separation (*a*), drift rate (*v*), gain sensitivity (*v*_*G*_), loss sensitivity (*v*_*L*_), and starting point bias (*z*), with 95% highest density intervals. **d**, Model comparison for depression symptoms using deviance information criterion (DIC). The value-sensitivity model (GL-mv) outperformed the baseline behavioural model (GL-m0), and was slightly better than the starting-bias model (GL-mz) and the boundary model (GL-ma). **e**, Posterior predictive check for accepted versus rejected reaction-time distributions from the GL-mv model with PHQ scores. **f-g**, Posterior distributions for Study II (Gain-Loss condition) with the same parameterization.

If affective symptoms alter the value-to-evidence interface – the stage where value information enters the accumulation process – effects should primarily appear in drift and its gain and loss components. By contrast, if symptoms primarily reflect altered decision policies, effects should manifest in boundary separation (a) or starting point bias (z). Within the DDM framework, threat avoidance predicts increased loss drift (*v*_*L*_ *↑*), reduced gain drift (*v*_*G*_ *↓* or unchanged), a starting point bias toward rejection (*z ↑*, biased toward rejection), and elevated decision threshold (*a ↑*) that slows commitment [2, 18, 24]. In contrast, reduced-motivation accounts predict reduced gain drift (*v*_*G*_ *↓*) and, in aversive contexts, reduced loss drift (*v*_*L*_ *↓*) and unchanged or reduced decision thresholds consistent with blunted motivational drive rather than increased strategic caution [19, 20]. Anxiety has also been linked to increased internal noise, which would manifest as less efficient evidence utilization without necessarily altering thresholds or starting biases [2]. Together, these predictions generate distinct computational signatures, allowing the competing theories to be tested within the DDM framework.

However, behavioural modelling alone might not fully determine whether apparent drift differences arise from degraded value encoding, altered accumulation dynamics, or ancillary systems such as attention and arousal [25]. Neural signals that track the evolving decision variable can provide additional constraints and improve the interpretability of behavioural models [26]. One such signal is the centroparietal positivity (CPP), a response-aligned, domain-general component that ramps up with accumulating evidence, predicts reaction time, and approaches a stereotyped amplitude at response – consistent with a stage linking valuation or sensory encoding to motor commitment [27–29]. Leveraging these properties, we implemented a neuro-informed hierarchical DDM in which trial-by-trial CPP features constrain specific parameters (slope *→* drift, peak *→* boundary, onset *→* non-decision time). This joint modelling improves parameter identifiability and allows a more direct test of where symptom-related distortions arise within the decision process.

In the present study, we used this approach to test whether anxiety and depression influence risky decision making primarily through heightened threat sensitivity and decision caution, or through diminished responsiveness to rewards. Across two culturally and economically distinct cohorts—a large behavioural sample from the United States and a behavioural–EEG sample from India—we instead observed a different pattern. Individuals with higher anxiety and depression scores exhibited reduced sensitivity to both gains and losses, reflected in weaker drift contributions from value, while decision thresholds and starting-point biases were largely unchanged. EEG analyses further showed that this diminished value sensitivity corresponded to the alterations in the dynamics of the centroparietal positivity (CPP), linking affective symptoms to the neural representation of accumulating evidence. Together, these findings suggest that anxiety and depression weaken the translation of value into evidence, rather than selectively amplifying threat avoidance or dampening reward pursuit.

## 2 Results

### 2.1 Choice Behaviour, Risk Aversion and Loss Aversion do not differ between anxious or depressed and the control groups

Participants completed a two-condition accept-reject gambling task designed to dissociate sensitivity to gains and losses (see Fig. 1a). In the Gain-Loss condition, each mixed gamble offered a 50% chance to win X and a 50% chance to lose Y, judged against a sure 0; in the Gain-Gain condition, gambles offered a 50% chance to win a high gain value and a 50% chance of winning a low gain value, again against a sure 100. Token values were later converted to monetary payment. This structure permitted separate estimation of gain- and loss-related influences on choice. Of the 201 participants in Study I, 62 met criteria for depression and 57 for anxiety, based on scores from the Patient Health Questionnaire (PHQ) (score *≥* 5) and Generalized Anxiety Disorder (GAD) scale (score *≥* 5), respectively [30, 31]. As reported in previous studies, PHQ and GAD scores were strongly correlated (r = 0.809, p *<* 0.001), resulting in substantial overlap between the depressed and anxious groups. Accordingly, behavioural outcomes across the two diagnostic categories were largely similar. These thresholded categories were used only for the simple descriptive comparisons below; all model-based analyses used continuous, z-score-normalized GAD or PHQ scores.

Overall, participants in the anxious and control groups exhibited comparable decision making patterns across task conditions. In the Gain-Loss condition, anxious participants accepted *M ± SD* = 29.9% *±* 23.3% of offers, slightly more than control participants, who accepted 25.4% *±* 23.8% of offers. However, this difference did not reach statistical significance (U = 3662.50, p = 0.090, Mann-Whitney U test). In the Gain-Gain condition, acceptance rates were nearly identical between anxious (26.6% *±* 24.7%) and control participants (26.9% *±* 23.4%; U = 4173.50, p = 0.853) (Fig. 1b). Similarly, response times also showed no difference (Fig. 1c), in the Gain-Loss condition, control participants responded in 1.1 *±* 0.4 (s), while anxious participants responded in 1.1 *±* 0.4 (s) (U = 4046.00, p = 0.877). In the Gain-Gain condition, control participants responded in 1.1 *±* 0.4 (s), while anxious participants responded in 1.1 *±* 0.5 (s) (U = 3787.00, p = 0.395).

Beyond acceptance rates, we estimated Prospect Theory parameters to assess effects on risk and loss aversion. Control participants showed a mean risk aversion of *M ±SD* = 0.5 *±* 0.1, while anxious participants exhibited slightly elevated estimates (0.6 *±* 0.7), though the difference was not statistically meaningful (U = 4091.00, p = 0.568) (Fig. 1f). Similarly, loss aversion parameters were nearly identical between control participants (1.65 *±* 0.76) and anxious individuals (1.70 *±* 0.83; U = 3988.00, p = 0.756) (Fig. 1e), suggesting comparable sensitivity to potential losses.

These findings replicated robustly in Study II (EEG study; Extended Data 1) and for depression (Extended Data 2), strengthening the conclusion that overt behavioural indicators of decision-making do not significantly differ between healthy and anxious or depressed individuals in this task. The absence of clear group-level differences in acceptance rates, risk preferences, or loss sensitivity suggests that affective disorders may not manifest in broad decision outcomes, but rather in more subtle, internal cognitive processes. Latent decision dynamics may therefore differ even when aggregate choice measures appear similar.

### 2.2 Value to evidence conversion is compromised in anxious and depressed

Given the absence of clear behavioural differences based on acceptance rates, we turned to a process-level analysis to investigate potential latent mechanisms underlying altered decision making. We fitted a hierarchical drift-diffusion model (DDM) (see Fig. 2a) to participants’ choices and response times. To formally test whether affective symptoms were associated primarily with altered decision caution, prior bias, or value sensitivity, we constructed a hierarchical family of models. We began with a baseline decision model (GL-m0), in which the drift rate (*v*) was driven solely by the trial-by-trial objective values of potential gains and losses. From this baseline, we developed competing extensions in which either anxiety or depression scores were included as population-level predictors of specific cognitive parameters.

Specifically, continuous symptom scores were modelled as modulating the decision boundary (GL-ma, testing the threat-avoidance hypothesis of increased caution), the starting point (GL-mz, testing for a prior bias toward rejection), or the drift rate and its gain/loss components (GL-mv, testing the value-attenuation hypothesis). This framework allowed us to directly estimate Bayesian regression parameters (*β*) across the model space to quantify the direction and magnitude of symptom effects.

Bayesian regression analyses across these model variants showed that continuous anxiety scores did not credibly affect boundary separation (*a*: *β* = 0.042, HDI = [-0.022, 0.107]) or starting point (*z* : *β* = -0.010, HDI = [-0.024, 0.004]). However, in the value-sensitivity model, loss sensitivity (*v*_*L*_) was reduced at higher anxiety scores, while gain sensitivity (*v*_*G*_) showed a weaker negative effect (*v*_*G*_: *β* = -0.014, HDI = [-0.027, 0.001]; *v*_*L*_: *β* = -0.019, HDI = [-0.032, -0.005]), with no direct symptom effect on overall drift rate (*v* : *β* = 0.025, HDI = [-0.096, 0.143]) (Fig. 2b). Thus, anxiety scores attenuated the contribution of losses to evidence accumulation, with a weaker and less certain effect on gain sensitivity, rather than biasing participants toward sampling accept or reject evidence overall. Continuous depression scores (PHQ) revealed a similar pattern: no credible effect on *a* or *z* (*a*: *β* = 0.026, HDI = [-0.038, 0.084]; *z* : *β* = -0.002, HDI = [-0.020, 0.013]), but reduced sensitivity to both gains and losses (*v*_*G*_: *β* = -0.023, HDI = [-0.037, -0.010]; *v*_*L*_: *β* = -0.014, HDI = [-0.028, 0.000]) (Fig. 2c).

Model comparison favored the value-sensitivity model for depression (GL-mv; DIC = 55371.76) over the baseline model (GL-m0; DIC = 55981.08) (Fig. 2d). The boundary and starting-bias models yielded nearby but higher values (GL-ma: DIC = 55378.28; GL-mz: DIC = 55378.53). For anxiety, the starting-bias, boundary, and value-sensitivity models had similar DIC values (GL-mz: 55378.23; GL-ma: 55383.04; GL-mv: 55383.47), so model comparison alone did not distinguish these regression structures. Posterior predictive checks showed that GL-mv reproduced the observed reaction-time distributions for accepted and rejected offers (Fig. 2e). These findings were replicated more strongly in Study II (Fig. 2f–g) (e.g., *v*_*G*_: *β* = -0.043, HDI = [-0.070, -0.016]; *v*_*L*_: *β* = -0.074, HDI = [-0.101, -0.047]), which matched our pre-registered hypothesis that both gain sensitivity (*v*_*G*_) and loss sensitivity (*v*_*L*_) would be reduced.

To test whether these disruptions in value sensitivity were specific to aversive contexts, we repeated the analysis in the Gain-Gain condition, where no losses were possible. In this condition, drift rate was estimated as a function of the high-gain and low-gain values (see Fig. 3a), so *v*_*G*_ and *v*_*L*_ index high-gain and low-gain sensitivity in this condition rather than gain and loss sensitivity. Continuous anxiety scores had no detectable influence on any DDM parameter (*a*: *β* = 0.028, HDI = [-0.042, 0.102]; *z*: *β* = -0.002, HDI = [-0.017, 0.012]; *v*_*G*_:GAD: *β* = -0.010, HDI = [-0.038, 0.018]; *v*_*L*_:GAD: *β* = 0.001, HDI = [-0.015, 0.018], all HDIs overlapping zero), suggesting that anxiety-linked gain insensitivity was contingent on the presence of loss (Fig. 3b). In contrast, continuous depression scores continued to predict reduced gain sensitivity (*v*_*G*_:PHQ: *β* = -0.041, HDI = [-0.066, -0.014]; *v*_*L*_:PHQ: *β* = -0.039, HDI = [-0.055, -0.023]). While *a* and *z* remained unaffected (*a*: *β* = 0.060, HDI = [-0.004, 0.124]; *z*: *β* = -0.005, HDI = [-0.017, 0.007]), consistent with a more stable reduction in reward sensitivity indicative of anhedonia (Fig. 3c). Model comparison showed that the value-sensitivity model (GG-mv) yielded a lower DIC (DIC = 51297) than the baseline model (GG-m0; DIC = 51315), indicating a better fit. The boundary model (GG-ma; DIC = 51317) and starting-bias model (GG-mz; DIC = 51303) had similar DIC values relative to the baseline model (Fig. 3d). Posterior predictive checks showed that the model reproduced the observed reaction-time distributions for accepted and rejected offers (Fig. 3e).

**Fig. 3.**
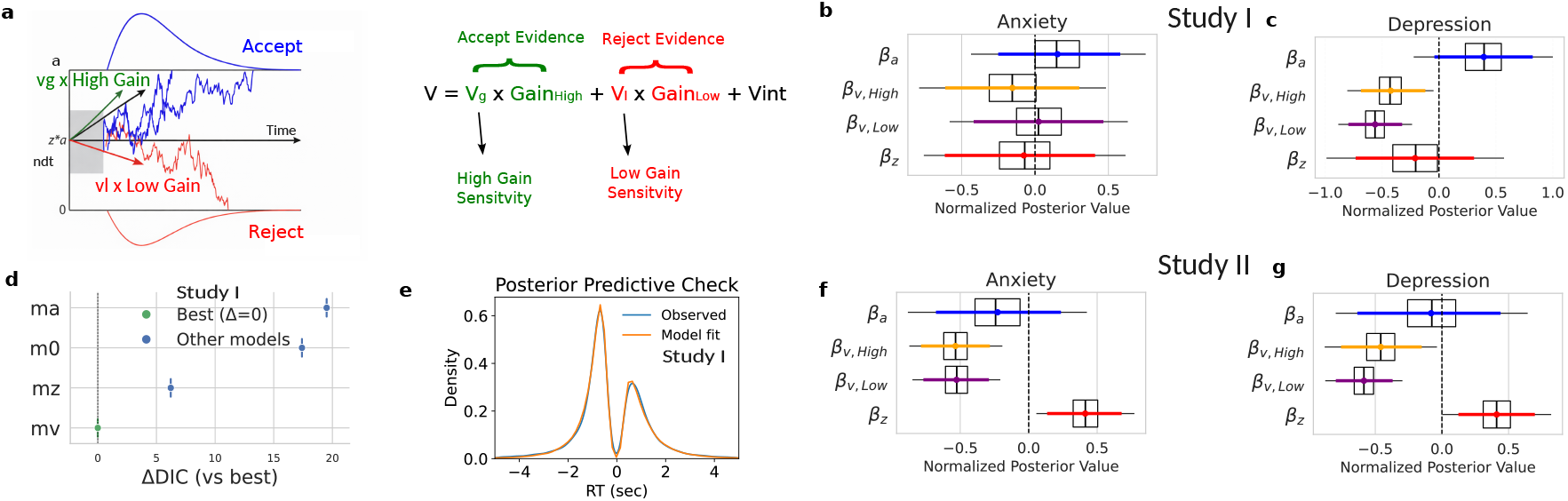
Drift diffusion modelling of Anxiety and Depression in the Gain-Gain condition. **a**, Schematic of the DDM in the Gain-Gain condition. Drift rate is defined as *v* = *v*_*G*_ ×*G*_*H*_ + *v*_*L*_ ×*G*_*L*_ + *v*_*int*_. **b-c**, Posterior distributions for regression coefficients on DDM parameters in Study I (Gain-Gain condition) as a function of continuous, z-score-normalized anxiety and depression scores. **d**, Model comparison for depression symptoms using DIC across GG-m0, GG-ma, GG-mz, and GG-mv. **e**, Posterior predictive check for accepted versus rejected reaction-time distributions from the GG-mv model with PHQ scores. **f-g**, Posterior distributions for Study II (Gain-Gain condition) with effects on *a, v, v*_*G*_, *v*_*L*_, and *z*.

This dissociation was not replicated in Study II. In the Gain-Gain condition, continuous anxiety and depression scores showed negative effects on the high- and low-gain sensitivity coefficients (Anxiety: *v*_*G*_:GAD: *β* = -0.109, HDI = [-0.159, -0.060]; *v*_*L*_:GAD: *β* = -0.135, HDI = [-0.192, -0.075] (Fig. 3f); Depression: *v*_*G*_:PHQ: *β* = -0.074, HDI = [-0.122, -0.028]; *v*_*L*_:PHQ: *β* = -0.154, HDI = [-0.208, -0.100]), with no reliable effects on *a, z*, or the drift rate *v* (Fig. 3g). These results mirror the findings from the Gain-Loss block and suggest a more generalised disruption in value sensitivity across both diagnostic groups in this sample. This pattern contrasts with Study I, where reduced gain sensitivity (*v*_*G*_) was selective to depression and anxiety effects overlapped zero. A simple explanation could be limited separability of GAD or PHQ collinear symptom dimensions in the smaller Study II sample.

The next section introduces a neuro-informed DDM in which trialwise neural covariates guide the value to evidence mapping. If limited separability in the smaller dataset is the primary cause, the Study I dissociation (depression-specific *v*_*G*_ or *v*_*L*_ reduction) should re-emerge once neural covariates partition variance in drift; if the joint Anxiety/Depression effects persist, this would instead support a shared reduction in gain sensitivity across internalizing symptoms.

### 2.3 CPP-Constrained Neuro-Informed DDM Localises Symptom Effects to Value Sensitivity

To evaluate the neurobiological plausibility of our model-based findings, we utilized centroparietal positivity (CPP), an EEG event-related potential (ERP) previously associated with the accumulation of decision evidence. Figure 4a displays the stimulus-locked and Figure 4c the response-locked CPP waveforms for both Gain-Loss and Gain-Gain conditions. As shown by the scalp maps, activity was maximal over a centro-parietal region. Time-frequency analysis indicated prominent *β*- and gamma-band modulation (Fig. 4b,d). Furthermore, no statistical difference was seen between the Gain-Loss vs Gain-Gain blocks when comparing the 95% CI of the CPP amplitude across the two conditions. These results suggest that the CPP reflects a domain-general evidence accumulation process that is not differentially modulated by the presence of potential losses.

**Fig. 4.**
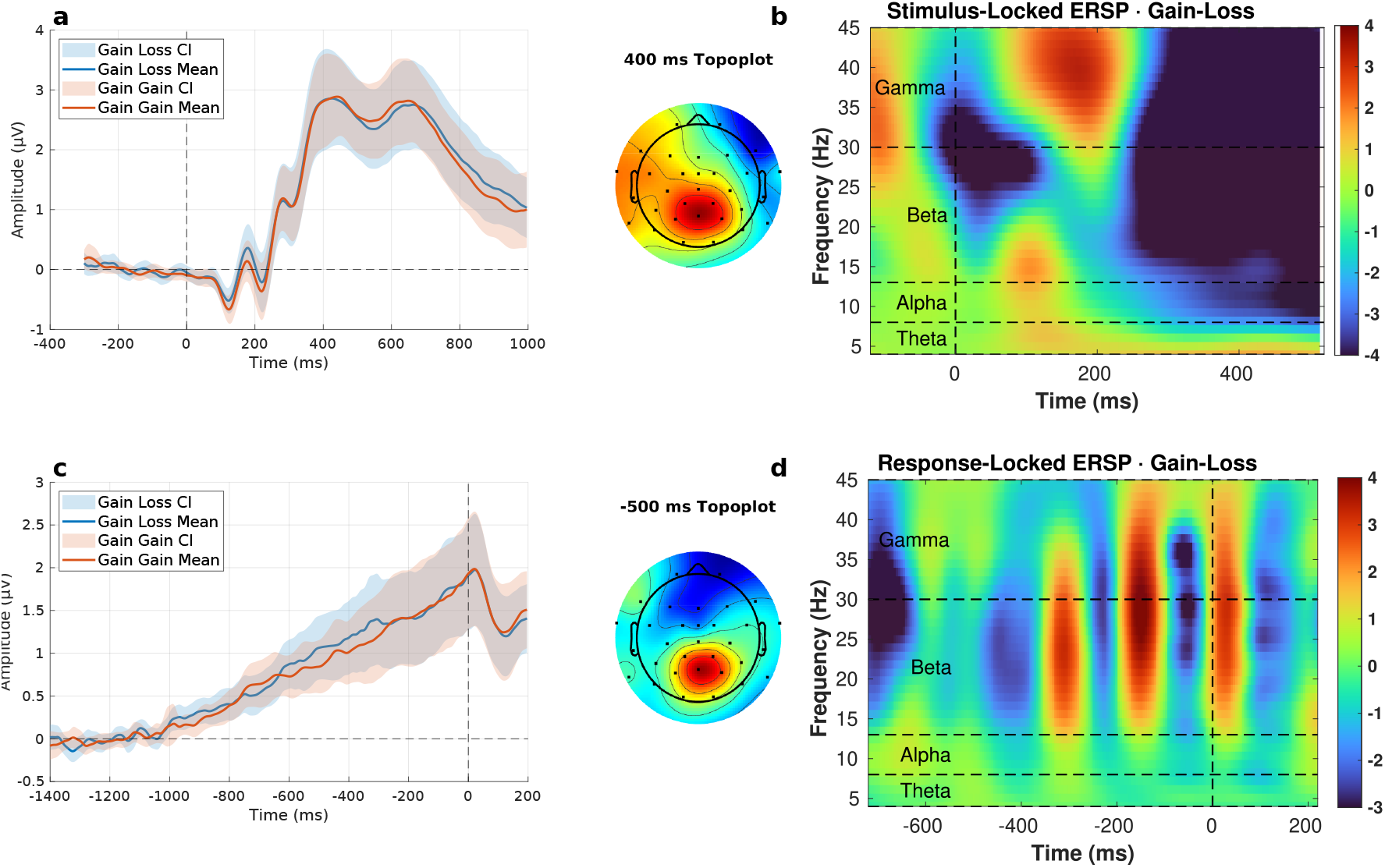
CPP dynamics in the gambling task. **a**, Stimulus-locked CPP at a centro-parietal region of interest (average of Cz, CP1, CP2 and Pz), shown for Gain-Loss and Gain-Gain conditions with 95% confidence intervals; scalp map at 400 ms. **b**, Stimulus-locked time-frequency representation from the same region. **c**, Response-locked CPP aligned to button press; scalp map at -500 ms relative to response. **d**, Response-locked time-frequency representation from the same region.

Previous studies have linked behavioural response-aligned CPP onset to non-decision time (*t*), CPP slope to evidence accumulation (*v*), and peak amplitude (CPP peak) to boundary separation (*a*) ([26, 28]). We extended this idea to the single-trial level using segmented regression (Fig. 5a), then we estimated onset latency, slope, and peak amplitude for each decision trial. To test the correspondence between EEG and model parameters, we used hierarchical Bayesian modelling to predict trial-wise DDM estimates from the CPP features. We compared several models: a behavioural-only DDM (EEG-m0), models using one EEG predictor each (EEG-mv, EEG-ma, EEG-mt), and a full combined model (EEG-mall). CPP peak significantly predicted boundary separation (*β* = 0.458, HDI = [0.417, 0.500]), and CPP slope significantly predicted drift rate (*β* = 0.046, HDI = [0.018, 0.072]). However, CPP onset did not reliably predict non-decision time (*β* = -0.003, HDI = [-0.008, 0.003]). The combined model (EEG-mall) had the lowest DIC (DIC = 6213.96), followed by the boundary-linked model (EEG-ma; DIC = 6368.54), while the behavioural-only model (EEG-m0) had the highest DIC (6918.17) (Fig. 5b). The Gain-Gain condition indicated similar results. These results indicate that CPP-derived neural markers significantly improve the precision of DDM parameter estimates and support the functional mapping of CPP features onto cognitive processes involved in value-based decision-making.

**Fig. 5.**
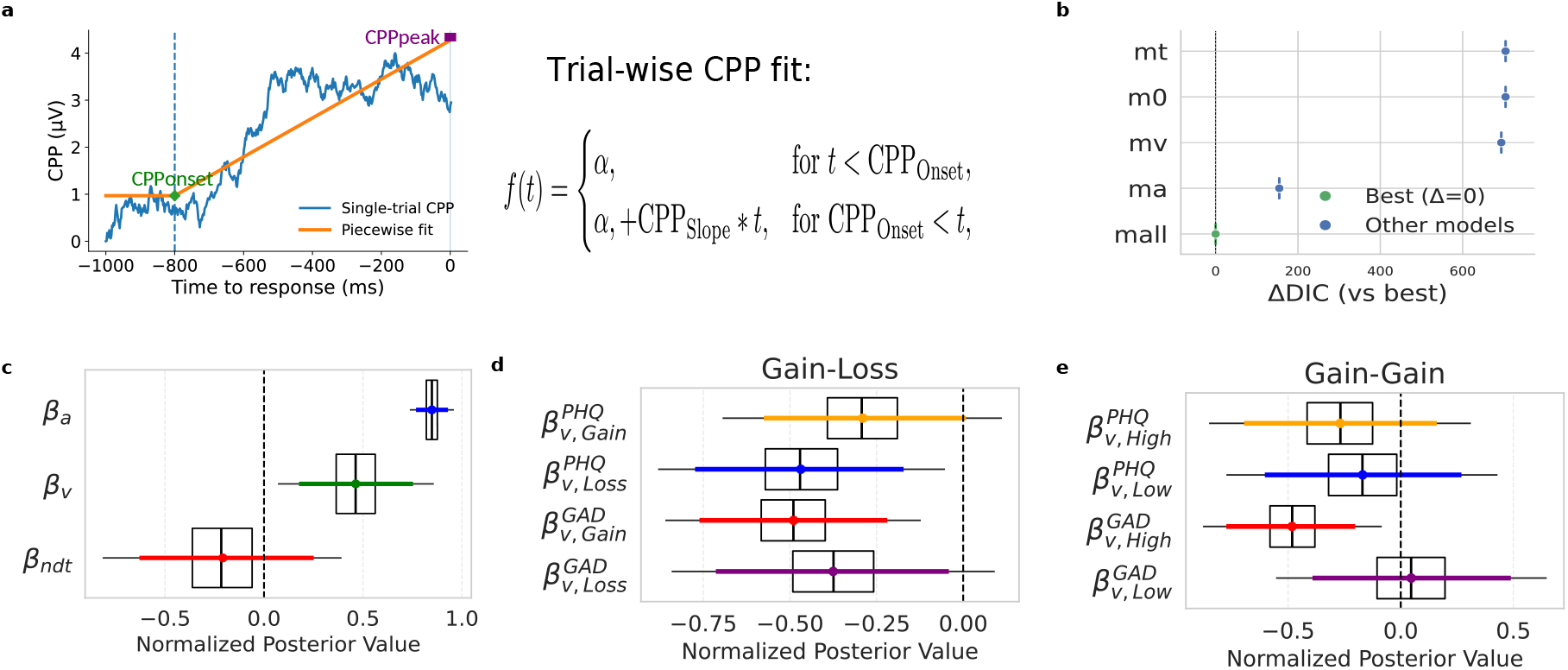
Linking CPP to DDM parameters and anxiety and depression symptoms. **a**, Single-trial response-locked CPP with piecewise linear fit used to extract onset time and post-onset slope. **b-c**, Model comparison (ΔDIC relative to best model) for the EEG family EEG-m0, EEG-mv, EEG-ma, EEG-mt, and EEG-mall in Gain-Gain and Gain-Loss datasets. **d**, Posterior summaries for the joint Gain-Loss models EEG-mav-GL-GAD and EEG-mav-GL-PHQ. **e**, Posterior summaries for the joint Gain-Gain models EEG-mav-GG-GAD and EEG-mav-GG-PHQ.

Re-estimating value sensitivity using the neuro-informed model in the Gain-Loss condition (Fig. 5d), continuous anxiety and depression scores predicted attenuated value sensitivity: anxiety reduced both gain and loss sensitivity (*v*_*G*_:GAD *β* = -0.064, HDI [-0.098, -0.030]; *v*_*L*_:GAD *β* = -0.040, HDI [-0.076, -0.006]), and depression similarly reduced *v*_*G*_ and *v*_*L*_ (*v*_*G*_:PHQ *β* = -0.036, HDI [-0.069, -0.001]; *v*_*L*_:PHQ *β* = -0.058, HDI [-0.092, -0.021]). However, in the Gain-Gain condition effects were weaker and more selective (Fig. 5e). The PHQ terms and the GAD effect on low-gain sensitivity overlapped zero (*v*_*G*_:PHQ *β* = -0.022, HDI [-0.053, 0.013]; *v*_*L*_:PHQ *β* = -0.015, HDI [-0.053, 0.020]; *v*_*L*_:GAD *β* = 0.004, HDI [-0.032, 0.040]), whereas the GAD effect on high-gain sensitivity remained negative (*v*_*G*_:GAD *β* = -0.057, HDI [-0.090, -0.026]). This pattern suggests that symptom-value disruptions are context-dependent and are weaker in the Gain-Gain condition (i.e., effects outside aversive context attenuate and often overlap zero). We therefore interpret the neuro-informed results as narrowing symptom effects to situations that include potential losses, with depression showing the most consistent, but context-amplified, reduction in gain-loss-related evidence.

Our preregistered hypotheses specified how neural decision signals would align with model parameters and how symptoms were expected to map onto the model parameters. H1 predicted that the trial-wise accumulation slope-indexed by the response-locked CPP would covary with the DDM drift rate (v) and decrease with greater anxiety/depression; this slope*→*drift mapping was supported and used to anchor v in the neuro-informed model. H2 predicted that the onset latency of the stimulus-locked CPP would track non-decision time (ndt); this mapping was not supported at the single-trial level, so ndt was left unconstrained by neural covariates. H3 predicted that the decision threshold (a) would scale with amplitude and increase with symptoms; while the CPP exhibited bound-like behaviour (peak amplitude relating to a), contrary to our hypothesis threshold was not affected by Anxiety or Depression. Incorporating CPP features improved fit and identifiability, thereby localising symptom influence primarily to value sensitivity (drift components) rather than to caution or prior bias, and setting up the interpretation developed in the Discussion.

## 3 Discussion

Across two complementary studies-a behavioural task and an EEG replication, we examined how anxiety and depression influence value-based decision-making. We tested two competing theoretical accounts: threat-avoidance models, which predict heightened sensitivity to potential losses and greater decisional caution, and reduced-motivation models, which propose blunted reward sensitivity and diminished responsiveness to gains.

Contrary to both predictions, our findings did not align neatly with either framework. Instead, symptom severity in anxiety and depression was associated with a *general reduction in value sensitivity*, rather than selective hyper- or hypo-sensitivity to only threat or reward. Hierarchical drift-diffusion modelling (DDM) revealed that higher GAD-7 and PHQ-9 scores predicted attenuated sensitivity to value information overall, while boundary separation (*a*) and starting point bias (*z*) remained largely unchanged. When drift rate (*v*) was decomposed into value-sensitive and value-independent components, symptom severity was associated primarily with reductions in the gain- and loss-dependent terms, while the value-independent component remained largely unchanged. This suggests that anxiety and depression weaken the influence of value information on evidence accumulation rather than globally slowing the accumulation process. Consequently, the primary behavioural signature of elevated symptoms is a weaker coupling between option value and the evolving decision variable, not necessarily slower responses overall. Because the task contains both attractive and unattractive gambles, value-dependent effects may partially cancel when reaction times are averaged across trials. Together with preserved boundary caution (*a*) and substantial trial-to-trial behavioural variability, this may explain why symptom-related differences are more apparent in latent decision parameters than in aggregate measures such as mean response times or acceptance rates. More broadly, these findings highlight the value of process-level computational models for detecting subtle alterations in decision-making that may remain hidden in conventional behavioural analyses.

The EEG study provided convergent neurophysiological support for this interpretation. The response-locked centroparietal positivity (CPP), a neural correlate of evidence accumulation ([27, 28]), tracked trial value and predicted behavioural responses. Incorporating CPP-derived parameters into the DDM significantly improved model fit (lower DIC) relative to behaviour-only models, and validated the reduction in value sensitivity.

In the Gain-Gain condition, depression-linked reductions in gain sensitivity replicated robustly, while anxiety effects were absent in the larger behavioural sample but evident in the smaller EEG cohort. When CPP-informed DDM models were applied, these effects also diminished in the Gain-Gain condition but persisted in the Gain-Loss condition, indicating that both disorders broadly attenuate value-to-evidence translation but did not selectively blunt reward pursuit or heighten loss sensitivity or caution. This pattern explains why acceptance rates appear unchanged even though model-derived parameters reveal a weakened mapping between objective value and the internal decision variable.

Taken together, these results suggest that anxiety and depression symptoms disrupt value-guided choice not through increased caution or directional bias, but through a broader dampening of sensitivity to both potential rewards and punishments. This domain-general reduction in value encoding points to a shared deficit in translating motivationally salient information into decision evidence, thereby providing a parsimonious account of how affective symptoms alter choice behaviour and its neural underpinnings.

Previous studies using mixed-gamble tasks have yielded inconsistent findings, with some reporting heightened risk sensitivity without stronger loss aversion in pathological anxiety [16], and others showing heterogeneous or null associations between affective symptoms and prospect-theoretic parameters [6]. One possibility for these inconsistencies could be reliance on descriptive model fits that capture choice patterns but overlook the latent processes that generate them. By decomposing decisions within a chronometric hierarchical drift-diffusion framework, our approach reconciles these disparate results. It shows that what appears as risk aversion at the behavioural level may, in fact, arise from a more fundamental reduction in value sensitivity-that is, weaker translation of objective gains and losses into evidence for choice. This process-level characterization reveals that anxiety and depression do not alter risk preferences per se, but rather diminish the fidelity of value signals driving evidence accumulation.

Embedding CPP features into the hierarchical model further improved predictive accuracy and reduced information criteria relative to behaviour-only baselines, demonstrating that neural signals impose non-redundant constraints on latent decision parameters. This brain-behaviour cross-validation strengthens the interpretation that affective symptoms impair the efficiency with which value information is transformed into decision evidence, rather than altering starting bias or decision thresholds.

Having localized this distortion to the value-to-evidence interface, we can now revisit the two prevalent theoretical accounts. Both threat-avoidance and reduced-motivation models capture partial aspects of the observed pattern, but neither fully explains the attenuation of value encoding seen here. Threat-avoidance accounts propose that anxiety heightens loss sensitivity and promotes cautious or avoidant decision policies, whereas reduced-motivation accounts emphasize blunted reward responsiveness and diminished approach drive. Our results diverge from both views, instead pointing to a shared impairment upstream of accumulation, likely at the value-to-evidence interface where affective symptoms reduce the fidelity of translating potential outcomes into decision evidence. Mechanistically, this disturbance suggests that interventions should focus on value representation rather than threshold control, by amplifying the salience and precision of goal-relevant value signals through affective forecasting training [32], attentional cueing of goal-relevant attributes, or reduced noise in value expectations. Instructions to attend closely to potential benefits, or visual highlighting of those benefits, may strengthen their representation during evidence accumulation. By contrast, exhortations to “be less cautious” are unlikely to remediate a deficit that does not principally involve boundary setting.

Several caveats merit consideration when interpreting these findings. First, anxiety and depression symptoms were moderately collinear, which constrains the separability of their individual contributions to reduced value sensitivity-particularly in the smaller EEG cohort. Although our analyses treated symptoms as continuous dimensions to preserve variance and avoid discontinuous diagnostic cutoffs, categorical overlap (e.g., GAD-7 and PHQ-9 *≥* 5) still limits the precision with which disorder-specific effects can be inferred. Future studies employing diagnostically characterized cohorts-primary anxiety, primary depression, and comorbid groups-will be essential to establish whether similar distortions in value-to-evidence translation are unique to either depression or anxiety.

Taken together, these findings provide convergent behavioural and neural evidence that anxiety and depression alter value-based decision-making not through increased caution or directional bias, but through a fundamental reduction in the efficiency with which value information is transformed into decision evidence. This value-to-evidence account reframes affective influences on choice as distortions in evidence quality rather than decision policy, integrating insights from computational modelling and electrophysiology. By localizing the dysfunction upstream of accumulation, at the interface where subjective value is encoded and translated into choice dynamics, the present work helps reconcile inconsistencies across prior descriptive models of risk and reward sensitivity. More broadly, these results underscore the utility of neuro-informed chronometric modelling for identifying mechanistic targets, suggesting that interventions aimed at enhancing the salience, precision, and stability of goal-relevant value signals may be more effective than those focused solely on modulating caution or control. In doing so, this framework advances a unified, process-level understanding of how affective symptoms disrupt motivated behaviour and decision-making.

## 4 Methods

### Gambling Task

Participants performed a simple gambling task (Fig. 1a). Each trial started with a 2–4 s gaze-fixation period, followed by presentation of a gamble featuring two equally likely outcomes (50%–50%), such as a gain of 20 or a loss of 60. Participants chose to accept the gamble, thus risking 60 for a chance to gain 20, or to reject it for a guaranteed alternative. These outcomes are in tokens, with an exchange rate of 20 tokens = $1. There were two trial conditions, the Gain-Loss condition, where one outcome was always Gain and the other always a Loss (e.g., + 50 or -60, with a guaranteed alternative of 0 if rejected), and a Gain-Gain condition, where both outcomes were gains (e.g. +150 or +40, with a guaranteed alternative of 100 if rejected). In the Gain-Loss condition, the guaranteed alternative was 0, while in the Gain-Gain condition it was 100. Each condition comprised 200 trials (100 unique gambles, each repeated once). These 200 trials were presented in 4 sets of 50 trials each, with a self-paced break between conditions. The order of the conditions and trials was randomised. Task performance was incentivised, and decisions directly affected participants’ payments. The show-up fee for the experiment was $15. At the end of the task, one trial was randomly selected from each condition and paid out in real money based on the participant’s decision during that trial.

### Data Collection

Data were collected in two phases, a behaviour-only Study I and an EEG Study II. Study I was conducted at the Wharton Behavioral Lab at the University of Pennsylvania, a total of 201 participants (mean age = 24.4 *±* 9.6 years, 138 females) volunteered. In Study II, participants were recruited at the Indian Institute of Technology, Kanpur, a total of 41 participants (mean age = 21.5 *±* 2.8 years, 13 females) volunteered. Both studies used closely matched tasks and designs. Study II included minor task changes for EEG recording; participants had a 5 s deadline to make their decisions, whereas Study I had no response-time limit. The token conversion was also changed from dollars to rupees to match the local currency. Study II was conducted after Study I data collection was complete. Study I was approved by the University of Pennsylvania Institutional Review Board (protocol number 827328), and Study II was approved by the Institute Ethics Committee of the Indian Institute of Technology Kanpur (reference number EEGfMRIDM20220818). All participants provided informed consent in accordance with the approved protocols. Standard self-report questionnaires were administered to quantify anxiety (Generalized Anxiety Disorder scale, State-Trait Anxiety Inventory) and depression (Patient Health Questionnaire, Beck Depression Inventory, and CES-D) [30, 31, 33–35]. These scores were used to examine the relationship between individual differences in affective symptoms and task behaviour.

### Pre-registration

Study II (EEG study) was conducted after Study I (Behavioural Study) data collection and was pre-registered based on preliminary Study I results. The pre-registration is available at AsPredicted (https://aspredicted.org/6k23-74j7.pdf) with reference ID #114276.

## Data and code availability

All data and analysis code are publicly available at https://github.com/mrugseniisc/GamblingEGGAnxDepp.

## Cognitive Modelling

Prior to modelling, trials with reaction times (RTs) faster than 200 ms or exceeding three standard deviations from the mean RT were excluded to eliminate anticipatory responses and outliers. Two modelling frameworks were then applied. First, Prospect Theory parameters, including loss aversion and risk aversion, were estimated within a hierarchical Bayesian framework implemented via the hBayesDM package [36] in R. Second, a hierarchical Drift Diffusion Model (DDM) was fit to both choices and RTs using the HDDM package in Python [37]. The DDM allowed trial-wise variation in drift rate.

In the Gain-Loss condition it was modelled as:

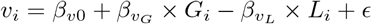

where *v*_*i*_ is the drift rate for trial *i, G*_*i*_ and *L*_*i*_ represent gain and loss magnitudes of the trial, *β*_*vG*_ and *β*_*vL*_ denote the respective hierarchical regression coefficients corresponding to gain-related and loss-related weight parameters, *β*_*v*0_ is an intercept, and *ϵ* is noise. In addition, we estimated hierarchical parameters for boundary separation *a*, starting point *z*, and non-decision time *t*.

In the Gain-Gain condition it was modelled as:

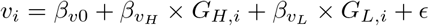

where *v*_*i*_ is the drift rate for the trial, *G*_*H,i*_ and *G*_*L,i*_ represent higher and lower gain magnitudes of the trial, *β*_*vH*_ and *β*_*vL*_ denote the respective hierarchical regression coefficients corresponding to high-gain-related and low-gain-related weight parameters, *β*_*v*0_ is an intercept, and *ϵ* is noise. In addition, we estimated hierarchical parameters for boundary separation *a*, starting point *z*, and non-decision time *t*.

### Effects of Anxiety and Depression

Effects of anxiety and depression were tested using GAD (Generalized Anxiety Disorder scale) and PHQ (Patient Health Questionnaire) scores. For descriptive summaries, participants were categorized as anxious or depressed if their GAD or PHQ score was *≥* 5, respectively. During model-based analyses, however, z-score-normalized GAD or PHQ values were used as continuous regressors to improve numerical stability and to avoid relying on hard diagnostic cutoffs. The thresholded labels were used only for descriptive summaries, and all inferential model comparisons used continuous symptom scores. For both conditions, we ran multiple hierarchical DDM variants that shared the same core latent parameters: drift rate (*v*), boundary separation (*a*), non-decision time (*t*), and starting bias (*z*). To avoid ambiguity across behavioural, EEG-linkage, and joint neuro-cognitive analyses, we refer to the models using the compact naming scheme in Table 1. The first prefix identifies the task or data family (Gain-Loss, Gain-Gain, or EEG), and the suffix identifies the parameter logic tested by the model.

**Table 1.** Model families used in the manuscript. HDDM regression equations map symptoms (GAD/PHQ) and/or EEG features to latent DDM parameters (*v, a, z, t*).

| Model Family | Model | Equation and Description |
| --- | --- | --- |
| Gain-Loss (GL) | GL-m0 | $v \sim \text{Gain} + \text{Loss}$<br>(Baseline reference; $v$ driven by objective values) |
| | GL-ma | $v \sim \text{Gain} + \text{Loss}, \quad a \sim \text{Symptom}$<br>(Tests threat avoidance/caution) |
| | GL-mz | $v \sim \text{Gain} + \text{Loss}, \quad z \sim \text{Symptom}$<br>(Tests prior bias toward rejection) |
| | GL-mv | $v \sim \text{Gain} + \text{Loss} + \text{Symptom} + \text{Gain} \times \text{Symptom} + \text{Loss} \times \text{Symptom}$<br>(Tests value sensitivity) |
| Gain-Gain (GG) | GG-m0 | $v \sim \text{Gain}_H + \text{Gain}_L$<br>(Baseline reference) |
| | GG-ma | $v \sim \text{Gain}_H + \text{Gain}_L, \quad a \sim \text{Symptom}$<br>(Tests caution in reward context) |
| | GG-mz | $v \sim \text{Gain}_H + \text{Gain}_L, \quad z \sim \text{Symptom}$<br>(Tests prior bias in reward context) |
| | GG-mv | $v \sim \text{Gain}_H + \text{Gain}_L + \text{Symptom} + \text{Gain}_H \times \text{Symptom} + \text{Gain}_L \times \text{Symptom}$<br>(Tests reward sensitivity) |
| EEG | EEG-m0 | (Behavioural baseline reference) |
| | EEG-mv | $v \sim \text{CPP}_{\text{slope}}$<br>(Links build-up rate to neural slope) |
| | EEG-ma | $a \sim \text{CPP}_{\text{peak}}$<br>(Links decision boundary to neural peak) |
| | EEG-mt | $t \sim \text{CPP}_{\text{onset}}$<br>(Links non-decision time to neural onset) |
| | EEG-mall | (Combines slope $\rightarrow v$, peak $\rightarrow a$, onset $\rightarrow t$) |
| Joint + Symptom | EEG-mav-GL | (EEG-ma + EEG-mv architecture + GL-mv regression structures for GAD or PHQ) |
|  | EEG-mav-GG | (EEG-ma + EEG-mv architecture + GG-mv regression structures for GAD or PHQ) |

#### Model structure

The behavioural family first splits into Gain-Loss and Gain-Gain branches, after which symptom regressors are added either to boundary separation, starting bias, or drift/value terms. The EEG family then adds CPP-derived covariates to the same DDM backbone and finally re-estimates the winning neural structure with symptom-linked drift terms.

### Behavioural Symptom-Model Family

Here, *v*_*G*_, *v*_*L*_, and *v*_*int*_ denote drift components, whereas *β* denotes hierarchical regression weights.

Across behavioural models, drift rate was modelled as a function of trial-level gain and loss magnitudes in the Gain-Loss condition:

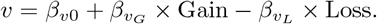

This formulation captures how participants integrated gains and losses into the decision process. The baseline behavioural model (GL-m0) estimated this value-driven drift function without any symptom-linked parameter modulation. The boundary model (GL-ma) added a symptom regressor to boundary separation,

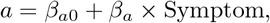

testing whether anxiety or depression altered decisional caution. The starting-bias model (GL-mz) instead placed the symptom regressor on

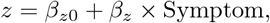

testing whether symptoms produced a pre-decisional response bias. The value-sensitivity model (GL-mv) allowed symptom scores to act directly on the value-to-evidence interface,

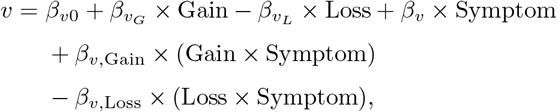

thereby testing whether symptoms altered how strongly gains and losses contributed to evidence accumulation. For depression analyses, the Symptom term was PHQ; for anxiety analyses, it was GAD. In the Gain-Gain condition, the same family structure was used with high-gain and low-gain trial values replacing gain and loss, yielding the parallel branch GG-m0, GG-ma, GG-mz, and GG-mv.

### EEG-Linkage Model Family

Within the EEG sample, we next asked which CPP features mapped most strongly onto which latent DDM parameters. The EEG baseline model (EEG-m0) omitted neural covariates. The drift-linkage model (EEG-mv) used CPP slope to predict drift rate, the boundary-linkage model (EEG-ma) used CPP peak amplitude to predict boundary separation, and the non-decision linkage model (EEG-mt) used CPP onset to predict non-decision time. The combined EEG model (EEG-mall) incorporated all three linkages simultaneously, allowing CPP slope, peak, and onset to jointly modulate *v, a*, and *t*.

### Joint Symptom + EEG Model Family

Finally, we fit joint symptom-plus-EEG models in which symptom-linked value sensitivity and trial-wise CPP covariates were estimated within the same hierarchical model. In these joint models, symptom terms entered value sensitivity, CPP slope entered drift, and CPP peak amplitude entered boundary separation. For the Gain-Loss condition, these were EEG-mav-GL-GAD and EEG-mav-GL-PHQ; for the Gain-Gain condition, these were EEG-mav-GG-GAD and EEG-mav-GG-PHQ. In each case, drift rate was modelled as a function of trial value, symptom score, their interaction terms, and CPP slope, while boundary separation was modelled as a function of CPP peak amplitude. This structure allowed us to test whether symptom-related attenuation of value-to-evidence translation persisted after explicitly accounting for neural signatures of accumulation.

Each model was fitted using hierarchical Bayesian inference with weakly informative priors using the HDDMregressor module of the HDDM package. Model comparison was based on the deviance information criterion (DIC). Regression coefficients were checked for highest density intervals (HDIs) overlapping zero to assess the presence or absence of reliable effects.

### EEG Recording and Preprocessing

Simultaneous electroencephalographic (EEG) data were recorded during task performance from a 32-channel system. Channels exhibiting abnormal standard deviations (using a threshold range of 1 to 100 SD) were removed and later reinstated by extrapolation, except when one of the critical CPP electrodes (CP1, CP2, Cz, Pz) was affected, in which case that participant was excluded. Seven participants were excluded on this basis, and one additional EEG file was unavailable because of recording failure. The remaining data were down-sampled to 250 Hz, band-pass filtered between 0.5 and 35 Hz, cleaned of 50 Hz line noise using CleanLine, and average-referenced. Independent component analysis was then performed, and components with *≥* 70% probability of representing eye or channel-noise artifacts were rejected using ICLabel [38]. A centroparietal potential (CPP) was derived by averaging signals from CP1, CP2, Cz, and Pz.

### Single-trial centroparietal positivity (CPP)

To characterize neural signatures of evidence accumulation on a fine temporal scale, we estimated singletrial centroparietal positivity (CPP) dynamics using segmented regression. Before fitting, the EEG signal was smoothed with a Gaussian kernel (standard deviation = 24 ms). Following Kraemer and Gluth [39], each CPP waveform was modelled with two connected linear segments: a pre-accumulation baseline and a rising segment leading to the response. Their intersection defined CPP onset, estimated by iteratively locating the breakpoint that minimized residual sum of squares. The slope of the rising segment indexed CPP build-up rate, providing a neural analogue of DDM drift. Each trial was fitted independently, yielding trial-wise CPP onset and slope estimates that could then be linked to latent decision parameters.

### Neuro-Cognitive Modelling

To formally link the neural and computational levels of analysis, we integrated single-trial CPP metrics into a hierarchical drift–diffusion modelling framework, extending beyond purely behavioural fits to construct a joint neuro-cognitive model of decision-making. This follows neurally informed and joint modelling approaches in which neural variables act as deterministic linking functions for latent parameters [40, 41], alongside work using EEG/fMRI covariates to predict trial-level decision parameters [42] and studies that constrain accumulation models with neural decision signals [26, 39], but here we explicitly incorporate trial-level EEG variability as continuous covariates in the DDM hierarchy. Individual estimates of CPP onset, slope, and amplitude were used to predict specific latent decision parameters, enabling us to test whether neural dynamics captured by the CPP track the same computational processes inferred from behaviour.

The EEG-linkage models therefore formed a nested family: EEG-m0 *⊂* {EEG-mv, EEG-ma, EEG-mt} *⊂* EEG-mall. For the joint symptom + EEG family, symptom terms acted on value sensitivity, CPP slope acted on drift, and CPP amplitude acted on boundary separation. For the Gain-Loss condition, drift rate took the form

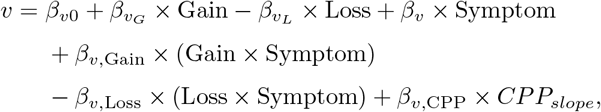

while boundary separation was modelled as

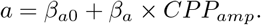

Starting bias (*z*) and non-decision time (*t*) were estimated hierarchically without additional neural covariates in these joint models. This structure allowed behavioural and neural predictors to jointly influence the rate of evidence accumulation, linking individual affective traits and neural decision dynamics within one model family.

### Model fitting

All models of both types, Prospect theory and Drift Diffusion were estimated hierarchically, i.e. the parameters were assessed at group level (i.e. for the entire subject population), as well at individual participant level (an estimate of deviation from the group values at participant level). Parameters were then estimated using MCMC sampling, in hBayesDM [36] for Prospect theory models and HDDM [37] for Drift Diffusion models. We ran 6 or 8 independent Markov chains for the models, depending on the size of the model. Each chain consisted of 3,000 to 4,000 samples, where the first 1,000 were burn-ins, yielding at least 2,000 posterior samples per chain. The models were assessed for convergence using the 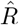 diagnostic; all 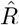 values were close to 1. Posterior predictive checks were also done on the best performing behavioural value-sensitivity model (GL-mv). Further GL-mv was also checked for parameter recoverability (Extended Data 3) and pairwise correlation between parameters (Extended Data 4). All these analyses showed that GL-mv converges well, its parameters are quite recoverable, and they do not trade off strongly with each other.

## Acknowledgements

We thank Wenjia Joyce Zhao for her earlier involvement with data collection and for providing feedback on the project. We also thank Peeusa, Avisha, and Shruti for support with data collection.

## Extended Data

**Extended Data 1: Behavioural results from Study II**

Behavioural results in Study I were qualitatively replicated in Study II. In the Gain-Loss condition, anxious participants accepted *M ± SD* = 34.4% *±* 17.0% of offers, compared with 31.9% *±* 13.7% in controls (U = 189.00, p = 0.786). In Gain-Gain, acceptance rates were also similar between anxious participants (49.0% *±* 20.7%) and controls (40.0% *±* 19.3%; U = 147.50, p = 0.163). Response times likewise did not differ reliably between groups. Fig. 6 summarizes acceptance-rate, response-time, and heat-map comparisons for Study II.

**Fig. 6.**
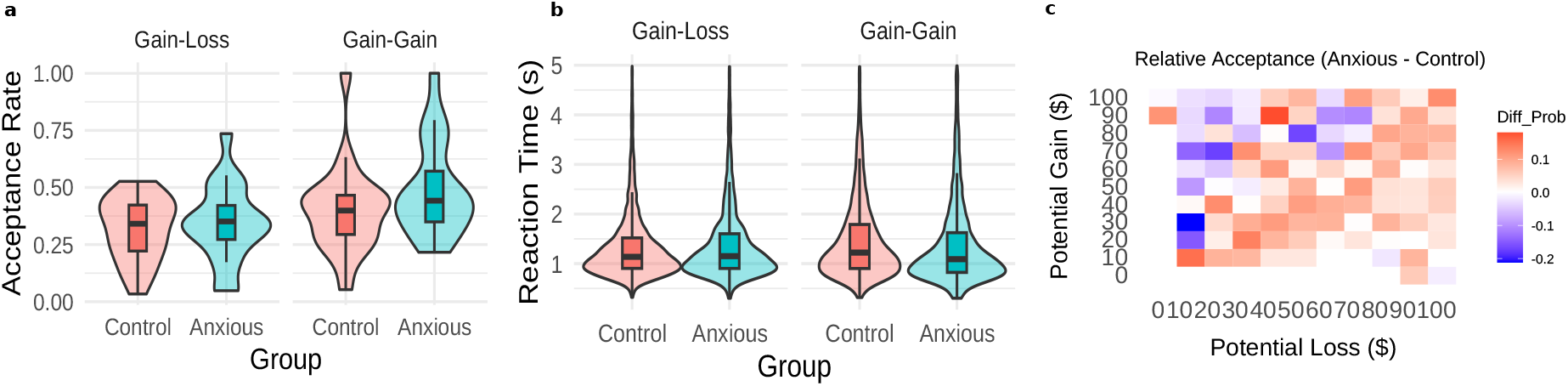
Behavioural results from Study II. Panel (a) shows acceptance rates as a function of gamble characteristics, panel (b) the corresponding response times, and panel (c) the heat map of group differences across gamble combinations. Behavioural differences between anxious and control participants were not statistically reliable.

**Extended Data 2: Behavioural results for depression**

Overall, depressed and control participants showed comparable behavioural performance across task conditions. In the Gain-Loss block of Study II, depressed participants accepted 33.6% *±* 16.3% of offers, compared with 31.0% *±* 11.1% in controls (U = 122.00, p = 0.582). In the Gain-Gain block, acceptance rates were likewise similar between depressed participants (45.5% *±* 19.3%) and controls (40.2% *±* 24.0%; U = 93.50, p = 0.141), and response times did not differ reliably in either condition. The same pattern was observed in Study I, where depressed participants accepted 29.9% *±* 20.9% of Gain-Loss offers versus 27.4% *±* 22.7% in controls (U = 3225.00, p = 0.263), and 42.5% *±* 27.9% of Gain-Gain offers versus 45.8% *±* 30.5% in controls (U = 3694.00, p = 0.768). Prospect Theory fits were similarly stable across groups in both samples: in Study I, loss aversion was 1.57 *±* 0.62 in controls versus 1.48 *±* 0.60 in depressed participants (U = 3920.00, p = 0.329), and risk sensitivity was 0.48 *±* 0.08 versus 0.47 *±* 0.09 (U = 3492.00, p = 0.755); in Study II, loss aversion was 1.59 *±* 0.53 versus 1.63 *±* 0.86 (U = 157.00, p = 0.588), and risk sensitivity was 0.86 *±* 0.14 versus 0.77 *±* 0.10 (U = 192.00, p = 0.092).

**Extended Data 3: Parameter recovery of GL-mv**

Using Study I posterior parameters as ground truth, we simulated 200 participants with 200 trials each and refit the model. Recovery was accurate for the primary drift components: *v*_*G*_ recovered at 0.502 [95% HDI 0.465, 0.538] against a true value of 0.480, and *v*_*L*_ at 0.633 [0.596, 0.669] against 0.600. PHQ terms were also well recovered and appropriately small: the PHQ main effect on drift was 0.022 [-0.012, 0.055] versus a true value of 0.023; the Gain *×* PHQ interaction was -0.037 [-0.074, 0.001] versus -0.023; and the Loss *×* PHQ interaction was -0.0068 [-0.043, 0.030] versus -0.014. Decision-non-drift parameters were tightly estimated as well, supporting identifiability of gain and loss sensitivity effects in drift (Fig. 7).

**Fig. 7.**
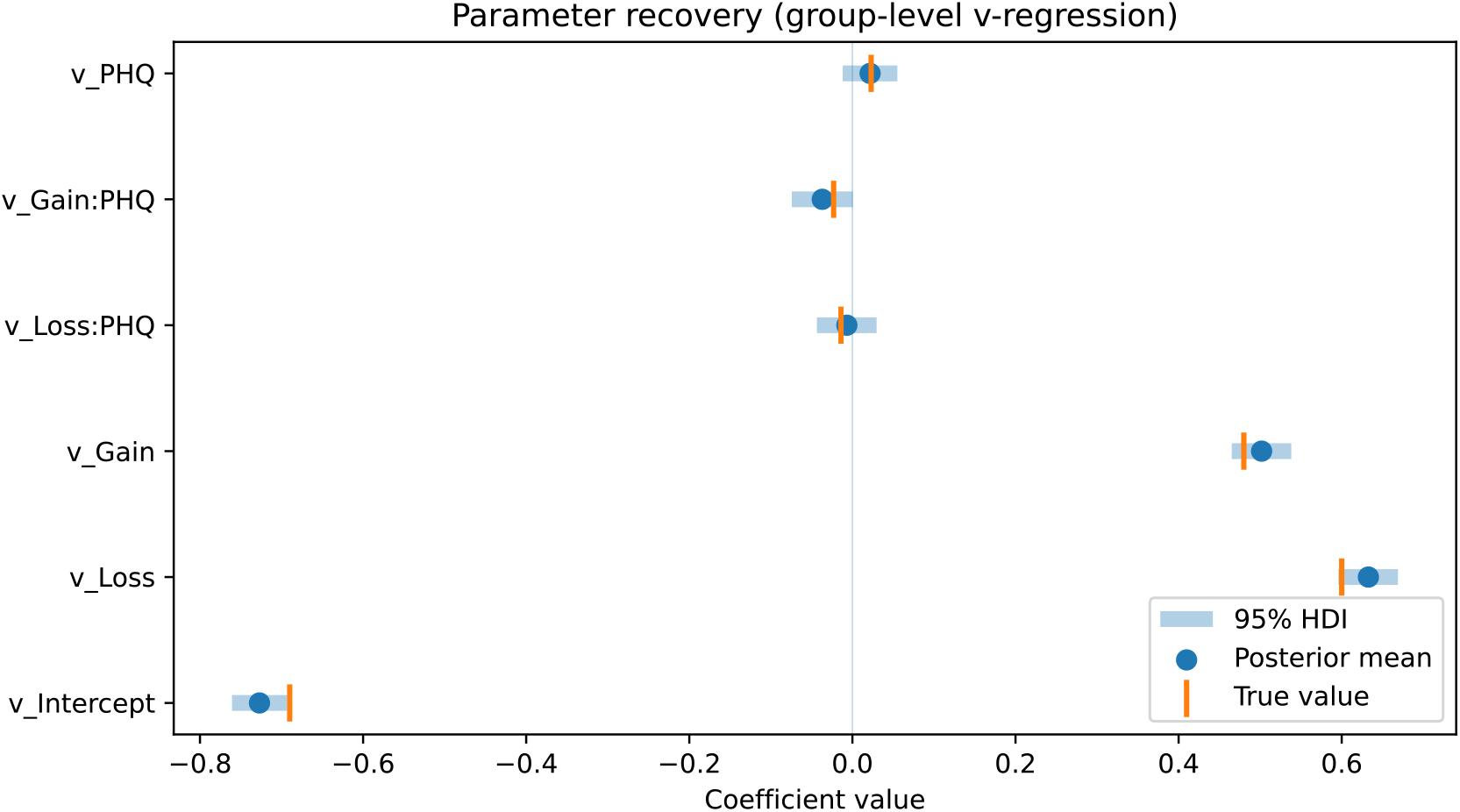
Parameter recovery for group-level coefficients in GL-mv. Posterior means (points) and 95% highest density intervals (bands) are shown for the drift intercept, gain and loss sensitivity terms, the PHQ main effect on drift, and the Gain× PHQ and Loss ×PHQ interactions. Orange ticks indicate the ground-truth coefficients used to generate the simulated dataset. Recovery was strongest for the primary value-sensitivity terms and remained close to the true values for the PHQ-linked coefficients.

**Extended Data 4: Pairwise parameter relationships**

Posterior draws showed compact, approximately spherical clouds for all parameter pairs involving *v*_*G*_, *v*_*L*_, the PHQ main effect on drift, and PHQ interactions with gain and loss. Drift sensitivities varied within narrow bands and did not co-vary strongly with PHQ terms, indicating limited collinearity and little evidence for compensatory parameter trade-offs (Fig. 8).

**Fig. 8.**
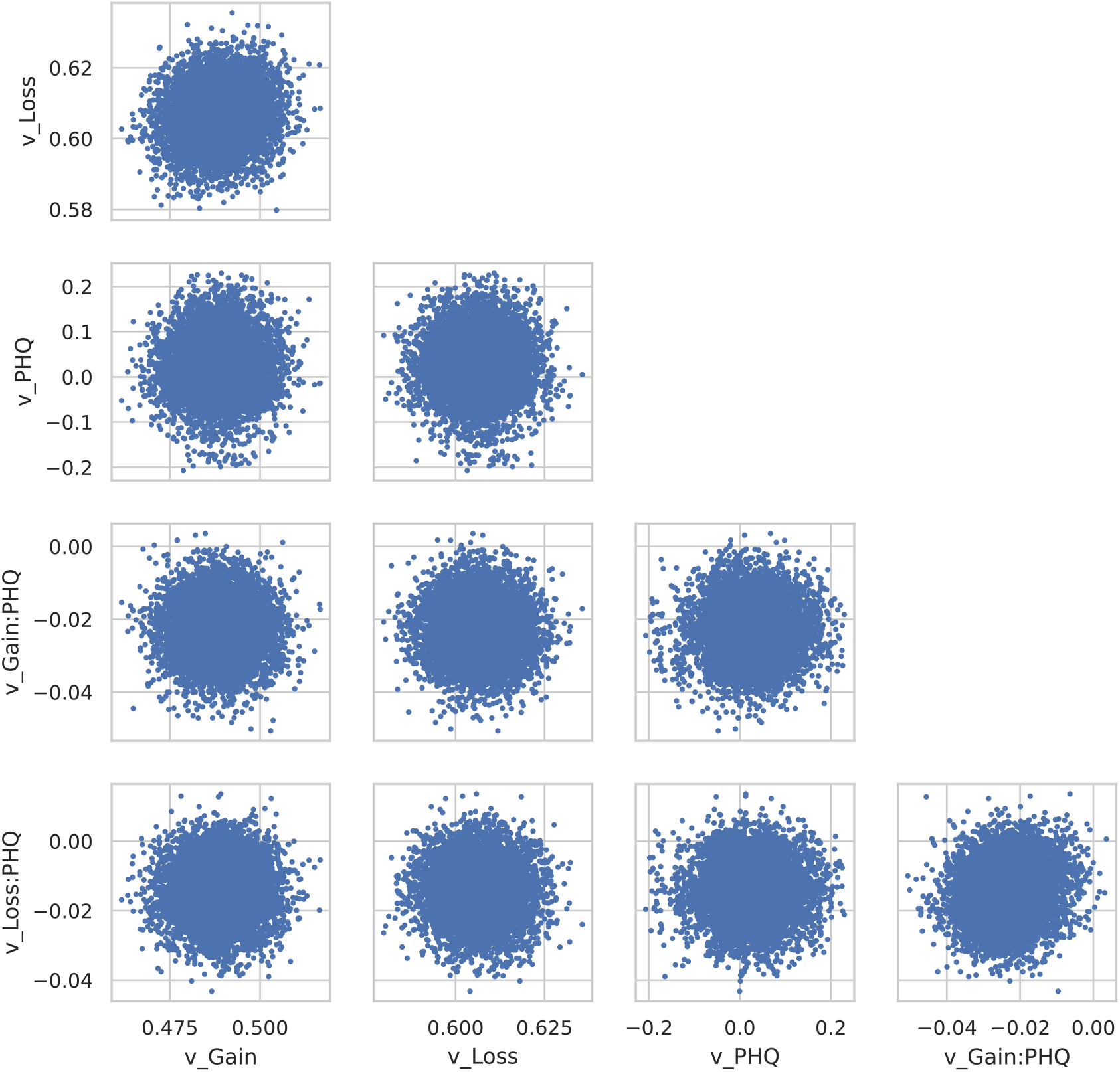
Pairwise relations among drift components and PHQ covariates. Scatterplots show joint posterior distributions for gain sensitivity (*v*_*G*_), loss sensitivity (*v*_*L*_), the PHQ main effect on drift, and the Gain× PHQ and Loss × PHQ interactions. Each point is one posterior draw. The clouds are compact and approximately circular, indicating limited pairwise dependence and little evidence for compensatory parameter trade-offs among the drift terms.

**Table 2.** EEG-DDM Bayesian parameter estimates for the Gain-Loss block. Posterior means, standard deviations, 94% highest density intervals, effective sample size for the posterior tails, and 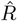 are shown for the principal EEG-DDM fits used in the Gain-Loss analyses.

| Model | Parameter | Mean | SD | HDI 3% | HDI 97% | ESS tail | $\hat{R}$ |
| --- | --- | --- | --- | --- | --- | --- | --- |
| EEG-m0 | $a$ | 1.905 | 0.071 | 1.771 | 2.039 | 8910.0 | 1.00 |
| EEG-m0 | $a_{std}$ | 0.382 | 0.059 | 0.280 | 0.495 | 8096.0 | 1.00 |
| EEG-m0 | $t$ | 0.581 | 0.026 | 0.531 | 0.629 | 11199.0 | 1.00 |
| EEG-m0 | $t_{std}$ | 0.147 | 0.022 | 0.107 | 0.185 | 8472.0 | 1.00 |
| EEG-m0 | $z$ | 0.506 | 0.012 | 0.484 | 0.530 | 9528.0 | 1.00 |
| EEG-m0 | $z_{std}$ | 0.235 | 0.024 | 0.191 | 0.279 | 8579.0 | 1.00 |
| EEG-m0 | $v_{int}$ | 1.534 | 0.119 | 1.313 | 1.758 | 8716.0 | 1.00 |
| EEG-m0 | $v_{int,std}$ | 0.593 | 0.085 | 0.443 | 0.755 | 10611.0 | 1.00 |
| EEG-m0 | $v_G$ | 1.151 | 0.040 | 1.075 | 1.223 | 4456.0 | 1.00 |
| EEG-m0 | $v_L$ | 1.427 | 0.041 | 1.349 | 1.503 | 3781.0 | 1.00 |
| EEG-ma | $t$ | 0.596 | 0.028 | 0.545 | 0.649 | 9830.0 | 1.00 |
| EEG-ma | $t_{std}$ | 0.155 | 0.022 | 0.115 | 0.196 | 8593.0 | 1.00 |
| EEG-ma | $z$ | 0.519 | 0.011 | 0.496 | 0.539 | 10405.0 | 1.00 |
| EEG-ma | $z_{std}$ | 0.213 | 0.023 | 0.169 | 0.257 | 8528.0 | 1.00 |
| EEG-ma | $v_{int}$ | 1.493 | 0.121 | 1.267 | 1.721 | 8197.0 | 1.00 |
| EEG-ma | $v_{int,std}$ | 0.593 | 0.086 | 0.445 | 0.756 | 10118.0 | 1.00 |
| EEG-ma | $v_G$ | 1.143 | 0.040 | 1.070 | 1.219 | 4481.0 | 1.00 |
| EEG-ma | $v_L$ | 1.409 | 0.042 | 1.332 | 1.491 | 3959.0 | 1.00 |
| EEG-ma | $a_{int}$ | 1.861 | 0.047 | 1.772 | 1.949 | 9880.0 | 1.00 |
| EEG-ma | $a_{int,std}$ | 0.240 | 0.039 | 0.172 | 0.315 | 8109.0 | 1.00 |
| EEG-ma | $a_{EEGa}$ | 0.458 | 0.022 | 0.417 | 0.500 | 10129.0 | 1.00 |
| EEG-mav-GL-PHQ | $t$ | 0.582 | 0.028 | 0.530 | 0.635 | 4905.0 | 1.00 |
| EEG-mav-GL-PHQ | $t_{std}$ | 0.156 | 0.023 | 0.115 | 0.199 | 3874.0 | 1.00 |
| EEG-mav-GL-PHQ | $z$ | 0.505 | 0.012 | 0.483 | 0.527 | 6139.0 | 1.00 |
| EEG-mav-GL-PHQ | $z_{std}$ | 0.234 | 0.023 | 0.193 | 0.278 | 4217.0 | 1.00 |
| EEG-mav-GL-PHQ | $v_{int}$ | -0.630 | 0.094 | -0.805 | -0.449 | 7370.0 | 1.00 |
| EEG-mav-GL-PHQ | $v_{int,std}$ | 0.516 | 0.074 | 0.384 | 0.653 | 4537.0 | 1.00 |
| EEG-mav-GL-PHQ | $v_G$ | 0.506 | 0.017 | 0.473 | 0.537 | 5330.0 | 1.00 |
| EEG-mav-GL-PHQ | $v_L$ | 0.651 | 0.018 | 0.619 | 0.686 | 5639.0 | 1.00 |
| EEG-mav-GL-PHQ | $v_G:PHQ$ | -0.036 | 0.018 | -0.069 | -0.001 | 6041.0 | 1.00 |
| EEG-mav-GL-PHQ | $v_L:PHQ$ | -0.058 | 0.019 | -0.092 | -0.021 | 5908.0 | 1.00 |
| EEG-mav-GL-PHQ | $v_{EEGv}$ | 0.167 | 0.015 | 0.138 | 0.194 | 5591.0 | 1.00 |
| EEG-mav-GL-PHQ | $a_{int}$ | 2.073 | 0.070 | 1.943 | 2.206 | 4737.0 | 1.00 |
| EEG-mav-GL-PHQ | $a_{int,std}$ | 0.369 | 0.055 | 0.274 | 0.474 | 4941.0 | 1.00 |
| EEG-mav-GL-PHQ | $a_{EEGa}$ | 0.429 | 0.024 | 0.386 | 0.474 | 3890.0 | 1.00 |
| EEG-mav-GL-GAD | $v_G:GAD$ | -0.064 | 0.018 | -0.098 | -0.030 | 6066.0 | 1.00 |
| EEG-mav-GL-GAD | $v_L:GAD$ | -0.040 | 0.018 | -0.076 | -0.006 | 6088.0 | 1.00 |

**Extended Data 5: EEG-DDM model output summary**

Selected EEG-DDM output summaries supported the main-text interpretation. In the Gain-Loss block, the EEG-informed models estimated positive couplings between CPP features and DDM parameters (*v*_*EEGv*_ = 0.167, HDI [0.138, 0.194], and *a*_*EEGa*_ = 0.429, HDI [0.386, 0.474], in the PHQ model), while symptom-related attenuation remained evident for both anxiety (*v*_*G*_:GAD = -0.064, HDI [-0.098, -0.030]; *v*_*L*_:GAD = -0.040, HDI [-0.076, -0.006]) and depression (*v*_*G*_:PHQ = -0.036, HDI [-0.069, -0.001]; *v*_*L*_:PHQ = -0.058, HDI [-0.092, -0.021]).

